# Newborn chicks prefer stimuli that move against gravity

**DOI:** 10.1101/2022.07.13.499929

**Authors:** Larry Bliss, Vera Vasas, Laura Freeland, Robyn Roach, Elisa Raffaella Ferrè, Elisabetta Versace

**Author notes:** Corresponding author; Elisabetta Versace, **Email:**. These authors contributed equally (Larry Bliss, Vera Vasas). **Author Contributions:** E.V. and E.F. conceived the research; L.F., R.R., L.B. and V.V. performed the research; L.B., V.V., L.F. and E.V. analyzed data; all authors wrote the paper.

## Abstract

At the beginning of life, inexperienced animals use evolutionary-given preferences (predispositions) to decide what stimuli attend and approach. Stimuli that contain cues of animacy, such as face-like stimuli, biological motion and changes in speed, are particularly attractive across vertebrate taxa. A strong cue of animacy is upward movement against terrestrial gravity, because only animate objects consistently move upward. To test whether upward movement is spontaneously considered attractive already at birth, we tested the early preferences of dark-hatched chicks (*Gallus gallus*) for upward vs downward moving visual stimuli. We found that, without any previous visual experience, chicks consistently exhibited a preference to approach upward moving stimuli, that move against gravity. A control experiment showed that these preferences are not driven by avoidance of downward stimuli. These results show that newborn animals are spontaneously attracted by upward movement, indicating that movement against gravity can be used as a cue of animacy to orient early approach responses in the absence of previous visual experience.

## Introduction

From the earliest stages of life, the ability to detect living beings is crucial for survival: living beings can give support (e.g., a mother hen provides warmth, protection and resources to chicks) or pose a threat (e.g., an approaching predator). In fact, early predispositions to approach/attend to cues of animacy have been found in young and inexperienced infants and other animals at the beginning of life (1). For instance, soon after birth, chicks, infants and tortoises preferentially attend to face-like stimuli (2–5), chicks and infants prefer objects that move in accordance with biological motion (6, 7), change their speed (8) or move along the longer body-plan axis (9, 10), as animate objects do. Movement against terrestrial gravity could provide important information on the presence of a living being because only animate objects can consistently move upward on their own. Accordingly, human adults find that upward moving stimuli appear more animate than downward moving stimuli (11). However, it is not known whether the association of upward movement and animacy is shared with nonhuman animals. It remains an open question whether animals require experience to associate upward trajectories with animate objects, or whether upward movements are attractive from birth, as an evolutionary-given preference.

To clarify whether newborn animals can use the upward direction of movement as a cue of animacy, we investigated whether newly hatched, visually inexperienced chicks (*Gallus gallus*) are spontaneously attracted by movement against gravity. Chicks are born with a mature sensory and motor system and have a strong motivation to approach social partners (1). Soon after hatching in complete darkness, with no experience with moving objects, we tested neonates’ preference to approach an upward moving vs a downward moving stimulus (a red circle attractive for chicks), presented on two opposite monitors for 20 minutes, divided in four consecutive time bins (Fig. 1A). Using automated tracking (12), we analysed the percentage of time spent close to the stimulus moving against gravity in each time bin, and the first-approach latency (time before entering a stimulus area).

**Fig. 1.**
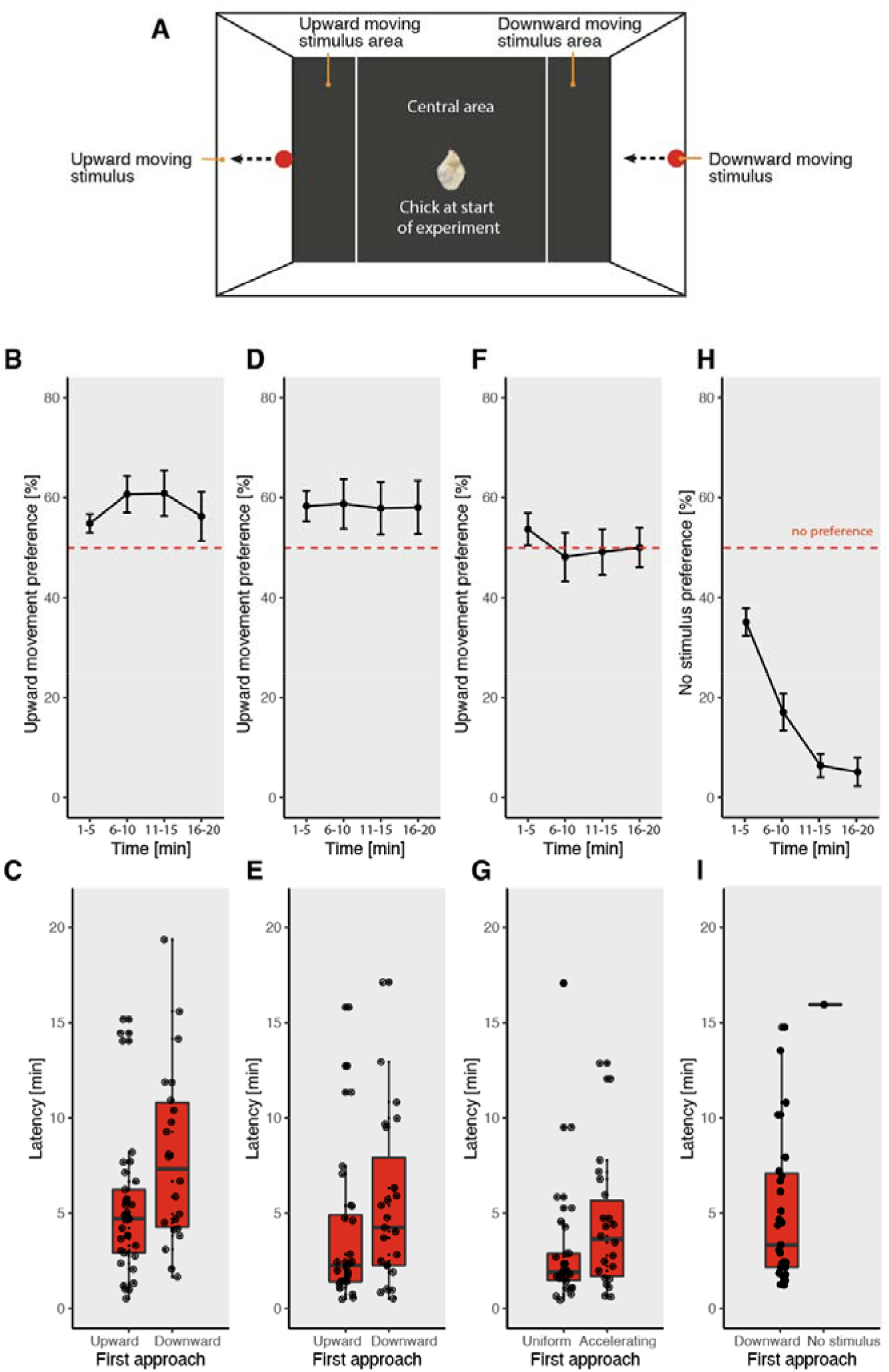
Chicks prefer upward moving stimuli. (A) Apparatus. The arena was divided into two stimulus areas and a central area. The position of the chick was recorded. (B) Preference in each 5-minute time bin and (C) latency for upward vs downward moving, accelerating stimuli. Preference (D) and latency (E) for upward vs downward constant speed movement. (F) Preference and (G) latency for upward uniform vs upward accelerating movement; Preference (H) and (I) latency for downward accelerating stimulus vs no stimulus. Bar plots and line plots show mean +/- SEM of preference. Boxplots display median, quartiles and outliers.

## Results

### Chicks spontaneously approach upward moving stimuli

In Exp. 1, we tested the preference for upward vs downward movement, presenting chicks with red circles that moved according to terrestrial gravity acceleration (9.81 m/s^2^) in opposite vertical directions (Movie S1, S5, see (13)). Chicks significantly preferred the upward moving stimulus: median 60.5%, W=476, p=0.014 (Fig. 1B). The first-approach latency was shorter for chicks that moved towards the upward stimulus (4.70 min vs 7.33 min, U=230, p=0.011, Fig. 1F). These results show that upward moving stimuli are more attractive than downward moving stimuli. In Exp. 2, we tested the preference for upward vs downward uniform (constant speed) movement (Movie S2). Chicks preferred the upward moving stimuli (median: 63.2%, W=448, p=0.044 (Fig. 1C), showing that the mere upward direction of movement (without acceleration) is sufficient to elicit a spontaneous preference. Again, chicks had a shorter first-approach latency for upward moving stimuli (median: 2.25 vs 4.25 min; U=234, p=0.049), Fig. 1G. In Exp. 3, we directly compared the responses to accelerating vs uniformly moving upward stimuli of the same average speed (Movie S3). Chicks showed no preference, spending 48.5% of time (median) at the uniform moving stimulus (W=587, p=0.992) (Fig. 1D). The first-approach latency was faster for the uniform upward movement (median: 1.90 vs 3.63 min; U=206, p=0.050), Fig. 1H. Overall, our results show that upward accelerating and constant speed movements are attractive for visually inexperienced chicks.

### Chicks do not avoid downward stimuli

Chicks are predisposed to flee away from fast looming approaching stimuli (14). To test whether the preference that we observed for upward moving stimuli was a fleeing response to downward, approaching moving stimuli, in Exp. 4 we tested chicks’ preference for downward accelerating movement vs a blank screen with no stimuli (Movie S4, S6). Chicks exhibited a very strong preference for the downward stimulus (median at 1-5 min=38.8, median at 6-10 min=1.7, p<0.001; median at subsequent times=0, p<0.001). These results show that the downward moving stimulus was not aversive (Fig. 1E and 1I).

## Discussion

The ability to discriminate between animate and inanimate objects is adaptive from early life, enabling animals to identify relevant information, find support from conspecifics and avoid threats (1). Upward movement against gravity is a clear – yet neglected – cue of animacy, since in most situations inanimate objects either remain still or move along gravity (e.g., a falling rock). We found that young, visually inexperienced chicks are attracted by stimuli that move upward, against gravity. These findings indicate that chicks have a spontaneous, non-learned predisposition for upward moving stimuli, suggesting that upward moving stimuli are used as a cue of animacy. Control experiments showed that the preference for upward moving stimuli does not depend on the presence of acceleration. In fact, chicks prefer upward vs downward stimuli irrespective of acceleration patterns and respond even faster to uniform movement. Moreover, the preference for upward movement is not driven by aversion to downward stimuli, since chicks were attracted by downward moving stimuli, when the other option was the absence of stimuli. Crucially, as chicks were completely visually naïve, our results show that the preference for upward movement is present in the absence of previous visual experience. These results pave the way to further studies on whether other species, including humans, are spontaneously attracted by upwards movement from early stages of life and on what dynamics and trajectories incompatible with passive movement along terrestrial gravity are attractive.

## Materials and Methods

At the end of incubation, chicks (*Gallus gallus*, 55 in Exp. 1, 51 in Exp. 2, 48 in Exp. 3, 31 in Exp. 4) hatched in darkness in individual compartments. Each chick was tested once for 20 minutes (four consecutive 5-minute bins) in a rectangular arena virtually divided in a starting area (54×60 cm), where the chick was initially located, and two stimulus areas (18×60 cm) (Fig. 1A). In the stimulus areas, two different video stimuli were simultaneously played in a loop on computer monitors (Asus MG248, 120 Hz), with left/right side counterbalanced between subjects. In Exp. 1, a red circle (Ø 3.28 cm), moved vertically upward or downward along the centre, with 9.81 m/s^2^ acceleration (Movie S1). In Exp. 2, the same stimuli moved at uniform speed (3.57 m/s) (Movie S2). Exp. 3 used the accelerating upward stimulus from Exp. 1 and the uniform upward stimulus from Exp 2 (Movie S3). In Exp. 4, one stimulus moved downwards at the acceleration of terrestrial gravity on one screen, with the other screen blank (Movie S4). Chicks’ centroid position in the arena was automatically tracked with DeepLabCut (12) and used to identify the position of the chick. Individual chicks’ preference was calculated as

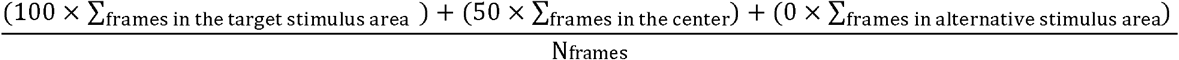

where 100 indicates a complete preference for the target stimulus, 0 for the alternative stimulus and 50 no preference.

Friedman tests showed no significant difference across the four 5-minute time bins in Exp. 1-3 (Exp. 1, Q=4.40, p=0.221; Exp. 2: Q=0.34, p=0.952; Exp. 3: Q=2.67, p=0.446). Thus, in Exp. 1-3 we calculated an overall preference index for each chick as an average of the four bins and tested the preference against chance level with a one-sample Wilcoxon test. The latency to first approach (first stimulus area entered) was assessed using a Mann-Whitney U test. In Exp. 4, the Friedman test detected a significant difference across time bins (Q=56.56, p<0.001), so the time bins were assessed individually with a Wilcoxon test. In Exp. 4, only one chick chose the No stimulus option, and for this reason differences in latencies could not be analysed. All tests were two-tailed, significance was set at p≤0.05. Analyses were run in Python, figures were prepared using R.

## Acknowledgments

This work was supported by the Leverhulme grant RPG-2020-287, the Royal Society Leverhulme Trust fellowship SRF\R1\21000155, and the BBSRC LIDO DTP programme.

## Supporting information

### S1 Extended methods

#### Subjects and Rearing Conditions

Experiments have been approved by the Queen Mary University of London ethics committee (AWERB) and Home Office (PP5180959). We used 193 domestic chicks, *Gallus gallus*, strain Ross 308. Overall, we tested 211 subjects: 61 chicks in Exp. 1 (2 excluded due to inaccurate tracking and 4 made no choice), 58 chicks in Exp. 2 (4 excluded due to inaccurate tracking and 3 made no choice), 59 chicks in Exp. 3 (11 made no choice), and 35 chicks in Exp. 4 (1 excluded due to inaccurate tracking and 2 made no choice). In Exp. 1, we analysed 17 males and 38 females (n=55), in Exp. 2, 16 and 35 (n=51), in Exp. 3, 22 and 26 (n=48), and in Exp. 4, 17 and 15 (n=32). All chicks were tested within 24 hours after hatching.

We incubated and hatched eggs in darkness in standard conditions (37.7 □C and 40-60% humidity) using a FIEM incubator S140ADS and hatchery H316DS). On day eighteen of incubation, eggs were moved to a dark hatchery in individual compartments.

#### Apparatus

The experimental arena (90×60×60 cm) was divided into three sectors for analysis: a central area (54×60 cm) and two stimulus areas (18×60 cm), see Fig. 1A. We recorded chicks’ performance with a camera located on top of the arena (resolution 1280×720 pixels, 10 fps).

#### Stimuli

The stimuli were displayed in an animation on two computer monitors (3840×1080 pixels at 120 fps). The animation was created with Unity 3D (2018, retrieved from https://unity.com/), and contained a red circle (Ø 3.28 cm) in the middle of a white screen. The average speed of the circle was 3.57 m/s. In Exp. 1 the circle was either moving downward or upward, according to terrestrial gravity acceleration (9.81 m/s^2^). In Exp. 2, the downward and upward stimuli moved at uniform (constant) speed. In Exp. 3, the circles were moving upward at accelerating (9.81 m/s^2^) vs uniformly speed. In Exp. 4, the downward moving stimulus presented in Exp. 1 was compared to no stimulus.

#### Experimental procedure

Immediately before the test, each chick was individually removed from the hatchery, in complete darkness, transported in an opaque box to the testing room (28 □C), sexed and checked for health. At the beginning of each 20 min test (four consecutive 5-minute time bins), video recordings started and the chick was gently placed in the middle of the arena, facing a long wall. After the test, chicks were housed in groups with food and water available ad libitum.

#### Video analyses

Videos were analysed through DeepLabCut (12), a markerless pose estimation system based on machine learning. We tracked chicks’ centroid position (as well as other markers on the chick’s body, to increase to precision of the centroid) and the three areas of the arena. To ensure high quality in behavioural tracking, for each marker we checked the likelihood tracking score of the markers, requiring a likelihood above 0.9 (this criterion was manually assessed as valid) for at least 90% of the frames. Seven subjects, whose tracking results did not reach this criterion, were not included in the analyses.

#### Data analyses

We analysed the preference for the target stimulus in each time bin and overall, and the first-approach latency (time elapsed until the centroid of the chick entered a stimulus area). The preference for the target stimulus was calculated as

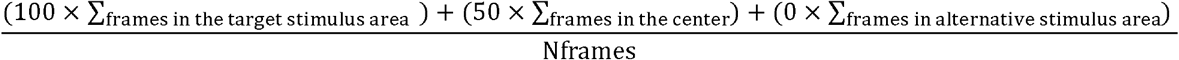

where 100 indicates a complete preference for the target stimulus, 0 a complete preference for the alternative stimulus and 50 indicates no preference.

Since data did not meet requirements for parametric statistics, we used the Friedman test to analyse the preference for the target stimulus across time bins, the one-sample Wilcoxon signed-rank test to identify significant deviations from chance level (50%) and the Mann Whitney test to check for significant differences in the latency between chicks that chose the target vs alternative stimulus. All tests were two-tailed. Significance was set at p≤0.05. Separate tests conducted with parametric statistics produced the same pattern of significance. Analyses were run in Python (libraries: numpy, pandas, scipy), figures were prepared using R (ggplot2).

